# American mammals susceptibility to dengue according to geographical, environmental and phylogenetic distances

**DOI:** 10.1101/2020.09.09.290031

**Authors:** Ángel L. Robles-Fernández, Diego Santiago-Alarcon, Andrés Lira-Noriega

**Affiliations:** Facultad de Física, Universidad Veracruzana, Xalapa, México; Red de Biología y Conservación de Vertebrados, Instituto de Ecología, A.C., Carretera antigua a Coatepec 351, Xalapa, C.P. 91070, México; CONACYT Research Fellow, Red de Estudios Molecualres Avanzados, Instituto de Ecología, A.C., Carretera antigua a Coatepec 351, Xalapa, C.P. 91070, México

**Keywords:** biotic interactions, DENV 1-4, machine learning, random forest, risk assessment, sylvatic cycle, wild reservoir

## Abstract

Many human emergent and re-emergent diseases have a sylvatic cycle. Yet, little effort has been put into discovering and modeling the wild mammal reservoirs of dengue (DENV), particularly in the Americas. Here, we show a species-level susceptibility prediction to dengue of wild mammals in the Americas as a function of the three most important biodiversity dimensions (ecological, geographical, and phylogenetic spaces), using machine learning protocols. Model predictions showed that different species of bats would be highly susceptible to DENV infections, where susceptibility mostly depended on phylogenetic relationships among hosts and their environmental requirement. Mammal species predicted as highly susceptible coincide with sets of species that have been reported infected in field studies, but it also suggests other species that have not been previously considered or that have been captured in low numbers. Also, the environment (i.e., the distance between the species’ optima in bioclimatic dimensions) in combination with geographic and phylogenetic distance is highly relevant in predicting susceptibility to DENV in wild mammals. Our results agree with previous modeling efforts indicating that temperature is an important factor determining DENV transmission, and provide novel insights regarding other relevant factors and the importance of considering wild reservoirs. This modeling framework will aid in the identification of potential DENV reservoirs for future surveillance efforts.

## 1 INTRODUCTION

Human impacts on natural environments have altered the structure and dynamics of ecological interactions (e.g., Maxwell et al. 2016), opening opportunities for a dramatic increase of novel emergent diseases (Jones-Engel et al. 2008; Allen et al. 2017). Given that viruses represent the main pathogenic group from where new emergent and re-emergent diseases affecting human populations come from, many efforts are directed toward viruses infecting mammals (e.g., Olival et al. 2017). Recent studies showed that emergent infectious disease events in human populations are more likely in regions with warmer and humid climates, places where land use change is directed toward agricultural systems and large urban areas, and in areas with higher mammal diversity (particularly rodents, bats, and non-human primates) (Allen et al. 2017; Olival et al. 2017; Gibb et al. 2020; Johnson et al. 2020; Mollentze and Streicker 2020). A disease that has increase its incidence in tropical regions due to human factors is the dengue virus (DENV), which is a mosquito-borne disease endemic to tropical areas and that has a sylvatic transmission cycle infecting bats and non-human primate species (e.g., Kato et al. 2013; Sotomayor-Bonilla et al. 2014). The natural sylvatic cycle has been altered due to anthropogenic activities, leading to an increase in its incidence for example by global warming that favors the geographic expansion of its mosquito vector (e.g., Brady and Hay 2020; but see Erickson et al. 2012 for a case where the dengue season is projected to be sorter due to the negative impact of climate change on the life span of the Asian tiger mosquito *Aedes albopictus*, an introduced species, in different cities of the USA). Thus, in order to be able to predict and prevent DENV expansion and increases in incidence, it is necessary to understand the ecological and biogeographic components of its sylvatic cycle (Vasilakis et al. 2011; Feldstein et al. 2015), particularly in the case of the American continent where the sylvatic enzootic cycle is poorly understood.

Studies on DENV have demonstrated that a large amount of variation in the incidence is explained by environmental variables such as temperature and precipitation, but there are no straightforward ways to generalize transmission dynamics given the complex interactions among climatic variables and local ecological factors (e.g., Morin et al. 2013; Morin et al. 2013; Feldstein et al. 2015; Messina et al. 2019). Furthermore, characteristics of a geographical range such as size and overlap (e.g., Olival et al. 2017; Dáttilo et al. 2020), and the richness of both vectors and host vertebrates are important determinants of a pathogen host breadth (Mollentze and Streicker 2020). In the case of DENV, the geographic expansion of one of its mosquito vectors (e.g., the invasive tiger mosquito) into places with immunologically naïve populations poses a major risk (Wearing et al. 2016; Brady and Hay 2020), particularly because researchers do not know ahead of time what species will enter the sylvatic cycle of the disease (e.g., Sotomayor-Bonilla et al. 2014). Thus, we need to develop more effective and preventive alternatives to reduce the risk of potential hazards becoming a real problem, and we also need to take the perspective of wild organisms and how they are affected by human activities (i.e., OneHealth or EcoHealth; Aguirre et al. 2012; Ostfeld and Keesing 2017). Ultimately we want to predict if the sylvatic cycle of DENV is geographically close to important human population sites, and how likely DENV is to switch among wildlife, domestic animals and humans (e.g., Gibb et al. 2020).

DENV is a pan-tropical self-limiting disease with a human death toll estimated in 10,000 fatalities and 100 million symptomatic cases per year (Messina et al. 2019). It is caused by a single-stranded RNA virus belonging to the *Flavivirus* genus, it is transmitted by different mosquito species (Diptera: *Culicidae*), and belongs to a viral family (Flaviviridae) that includes other major diseases such as yellow fever (YFV), West Nile (WNV), and Zika viruses (Brady and Hay 2020). DENV includes at least four genetically different serotypes or clades, which are embedded in human urban cycles (mostly transmission among humans via mosquitoes of the genus *Aedes*, in particular *Aedes aegypti*) and also in ecologically complex sylvatic cycles (Sotomayor-Bonilla et al. 2014; Brady and Hay 2020). Regarding sylvatic cycles, there are only two places where transmission cycles are reasonably well determined, one is in Senegal and the other is in Malaysia where non-human primates belong to a permanent sylvatic cycle (Vasilakis et al. 2011; Althouse et al. 2015). Some researchers suggest that sylvatic cycles are not established in the Americas because only human-adapted DENVs have been identified infecting wild animals (e.g., Morales et al. 2017). Yet, researchers modeling expansion of DENVs have done it via geographic projections of *Aedes* spp. vectors (e.g., Padilla-Torres et al. 2013; Butterworth et al. 2017), and little effort has been invested in determining the vertebrate mammal assemblages potentially maintaining DENV sylvatic cycles in the Americas. Moreover, every human DENV serotype has a sylvatic ancestor (Costa et al. 2012) and well-known mammal Orders (i.e., primates, rodents and bats) that are reservoirs for many viral families are readily infected with DENV in Latin America. Thus, many wild mammal species may belong to the enzootic sylvatic cycle of DENV in the Americas (Thoisy et al. 2009; Sotomayor-Bonilla et al. 2014; Morales et al. 2017), and it is a medical and veterinary priority to discover who those species are.

The use of analytical statistical tools can take advantage of knowledge on host breadth, environmental variables, geographic distributions, and host phylogenies to develop predictive frameworks to generate geographical assessments of infection risk, which can be applied to either host assemblages or a focal host species by either a group of pathogens or by a specific pathogen of interest (e.g., Robles-Fernández and Lira-Noriega 2017). Thus, our aim is to predict sylvatic host assemblages of DENV in the Americas via machine learning statistical protocols, in order to create preventive frameworks useful for health agencies (e.g., CDC) that can aid to forecast the probability or risk of potential hazards (Hosseini et al. 2017). Here, we analyze the relevance of geographic, environmental, and phylogenetic distances among potential wild host species of DENV to predict their susceptibility to the virus. This predictive framework allows to detect host assemblages or species with higher risk of infection across these three biodiversity dimensions, as well as to identify potential host species where the pathogen has not been previously detected or that have not been previously considered in field screening projects. This approach will help to direct surveillance and field efforts providing cost-effective decisions on where to invest limited resources to untangle DENV sylvatic cycles in the Americas, in case they are already established.

## MATERIALS AND METHODS

### Host -parasite data

We did an exhaustive review of information about the incidence of dengue virus in mammals. We started from the information available in Sotomayor-Bonilla et al. (2019), from which we obtained the references and subsequently obtained information from 15 articles. This search allowed us to collect a total of 249 incidences in 30 species of mammals distributed geographically in the Americas (Supplementary Material). Since only 10 species of mammals contained 80% of the information, we decided to use this as the initial susceptibility cutoff threshold to conduct model training (Figure 2). Thus, model predictions will be consistent with the environmental, geographic and phylogenetic information from this set of species.

**Figure 1.**
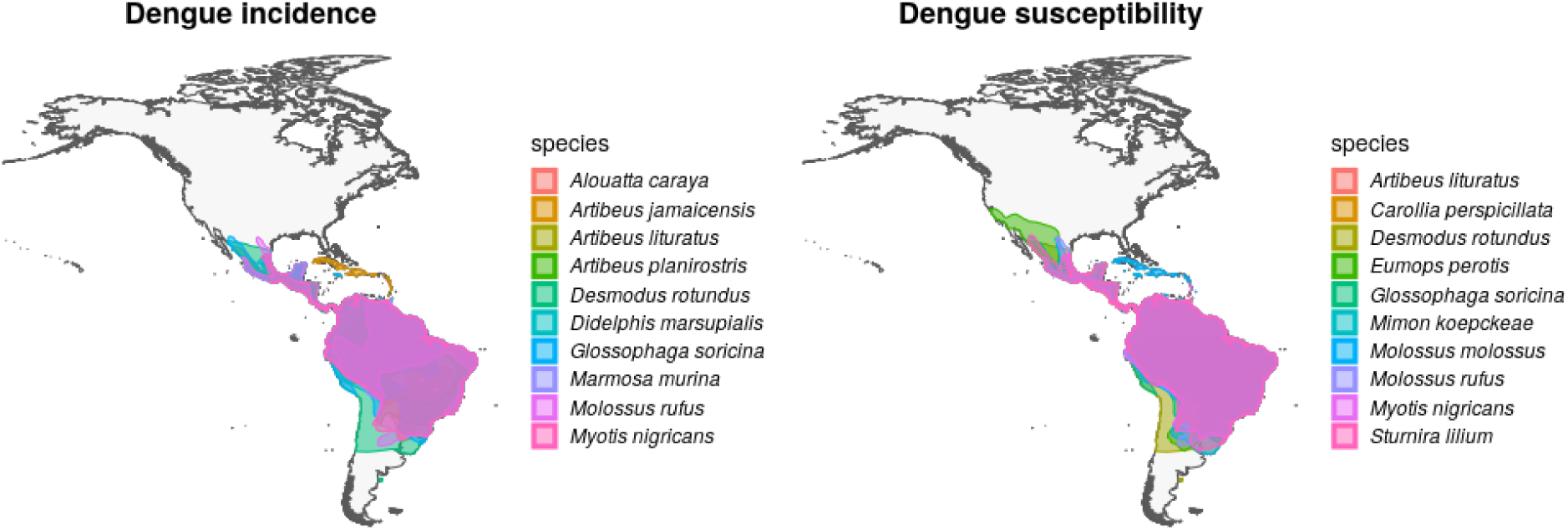
Geographic distribution of mammal species with high incidence and susceptibility to DENV. Left panel shows the top ten wild mammal species with highest DENV incidence according to published information. Right panel shows the top ten most DENV susceptible wild mammal species according to the random forest models.

**Figure 2.**
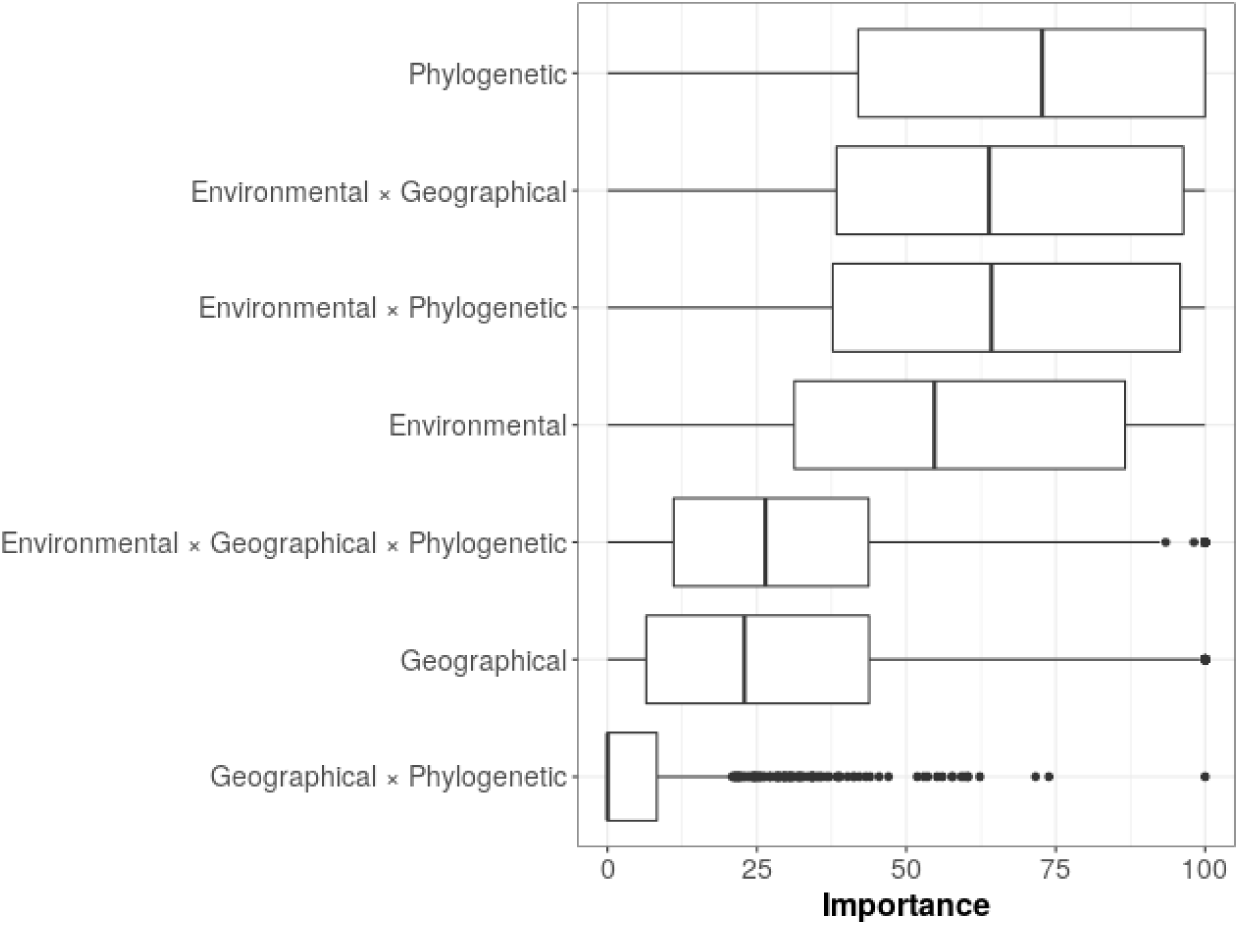
Variable importance from random forest models. The mean percentage values for the variable importance are as follows: phylogenetic = 67.5; environmental*geographical = 62.6; environmental*phylogenetic = 62.3; environmental = 55.8; environmental*geographical*phylogenetic = 29.9; geographical = 28.5; geographical*phylogenetic = 6.38.

### Input data processing

We estimated the susceptibility to DENV based on species’ distances from the geographic, environmental and phylogenetic spaces. To calculate the geographic distance matrix between pairs of species, we calculated the geographic distance between the centroids of the largest polygon for each mammal species in the Americas available from the International Union for Conservation of Nature polygons (International Union for Conservation of Nature 2020; the IUCN Red List of Threatened Species, version 2018-2. https://www.iucnredlist.org. Downloaded on 05 January 2020). A total of 1778 species of land mammals were selected for America. We carried out this geographical data manipulation with the R package sf (Pebesma 2018).

Then, for between-species environmental distance, we estimated the distance between the maxima of the probability density with a Gaussian kernel associated with the first two principal components from the 19 bioclimatic layers from the WorldClim database (approximately 80% of the total variance explained; Fick and Hijmans 2017). The probability density was based on a sample of 10000 points contained within the geographic range of the species based on the IUCN polygons mentioned above. In this case, the maxima of the probability density were considered as a surrogate of the ecological niche centroid for each species. The principal components were based on a correlation matrix on layers with a 2.5 minutes spatial resolution; the use of these layers was based on the idea of using a low dimensionality of environmental variables while accounting for most of the information. For phylogenetic distances, first we generated a pruned phylogenetic tree with the species of terrestrial mammals of the American continent, which we then used to calculate the phylogenetic distances between pairs of species with the ape R package (Paradis and Schliep 2019). The original phylogenetic tree was from Faurby et al. 2018. All of the above information was summarized in a database with the environmental, geographic and phylogenetic information among all pairs of species (∼3 000 000 rows). Next, we took the average of one species with respect to all others in order to summarize the information about the three distances (i.e., geographic, environmental, and phylogenetic). This rendered a table of 1778 species x three distances.

### Machine learning modelling

For each host-parasite assemblage, we created a data set taking each *n* host species and the phylogenetic, environmental, and geographic distances calculated above. Based on this data set, we selected the species considered in the pathogen incidence cutoff (see above) and labeled these species as susceptible. Subsequently, we took a random sample of species outside of the cutoff set, balancing the sample size with respect to the cutoff set and labeling these species as unknown. This generated a data set with two susceptibility classes (susceptible and unknown) and three independent variables (environmental, geographic and phylogenetic distances). To this data set, we also added the interactions between the independent variables and separated the data set into two sets to train the models and validate them (70% - 30%). Following Kuhn (Kuhn et al. 2008), we generated a set of random forest models optimizing their parameters with a modeling grid (Kjeldgaard 2018) and with 10 times 10-fold cross validation for each sample. With the optimal model we regressed the entire data set and obtained the probability of susceptibility given the distances and their interactions, also calculating the importance of each variable. Each time we classified the species as susceptible with a standard probability threshold (i. e. *p*(*x*) *>* 0.5). We repeated this procedure 1000 times, each with a different sample, in order to look for convergence in the probability calculation, as well as its uncertainty. We then estimated the mean probability of susceptibility for each host species in each host-pathogen assemblage, along with its standard error (Hastie et al. 2009).

### Susceptibility geographical richness

In order to provide a spatial pattern of susceptibility to dengue across American mammals, we assigned the average susceptibility probability of each host species to each of their geographic ranges. Subsequently, we selected only species with probability of susceptibility *p*(*x*) *>* 0.5 and considered these as susceptible. Finally the geographic ranges of susceptible species were used to generate a species richness map of potential dengue hosts on the geographic space. This was done using the raster (Hijmans 2020) and fasterize (Ross 2020) R packages.

### Statistical validation of spatial patterns

To test our model on geographic space, we placed empirical validation points of where dengue has been found in the field and applied a statistical hypothesis on the spatial distribution of the punctual empirical pattern with respect to our susceptibility richness map. We tested as null hypothesis (*H*_0_) that the density of empirical pathogen points is not a function of the richness of susceptible host species, and as an alternative hypothesis (*H*_*a*_) that the density of empirical pathogen points depends on the richness of susceptible host species according to the random forest model. We performed this test using the likelihood ratio test (LRT) for each hypothesis, getting in all cases *p <* 0.05 and rejecting *H*_0_ in all cases. We implemented this analysis with the R package spatstats (Baddeley et al. 2004; Gelfand et al. 2010).

## 2 RESULTS

After overlapping the ranges of 60 susceptible species (i.e., species with probability of susceptibility *p*(*x*) > 0.5), the species richness map for susceptible species to DENVs showed higher values in tropical America and decreasing values toward southern South America and western North America (Figure 3). This pattern was statistically corroborated by independent localities with incidence of DENV in mammals, indicating that it follows a spatial point pattern depending on species richness predicted as susceptible, with the highest density of DENV in mammals predicted at a species richness close to 20 (Figure 3). The mean susceptibility to DENV from the 1774 mammal species evaluated followed a similar geographic trend with respect to susceptible species richness (Figure 3 B). The top 10 species with highest susceptibility to dengue are shown in Table 2. These belong to bats of different trophic guilds, mainly insectivores, frugivores, and nectarivorous, but the species at the top of the list is the common vampire bat that feeds primarily on mammalian blood (Table 2). Interestingly, the pattern of DENV intensity given richness of susceptible species showed two regions with high values (Figure 4). One appears in central Mexico in the northern limit of the Neotropical biogeographic region, in what looks like a belt that connects the gulf of Mexico and the Pacific coast across continental Mexico, and an isolated portion at the north of the Yucatan Peninsula. The second region of high intensity appears in South America, also in what looks like a belt that connects the southern portion of Brazil to northern Peru (Figure 4).

**Figure 3.**
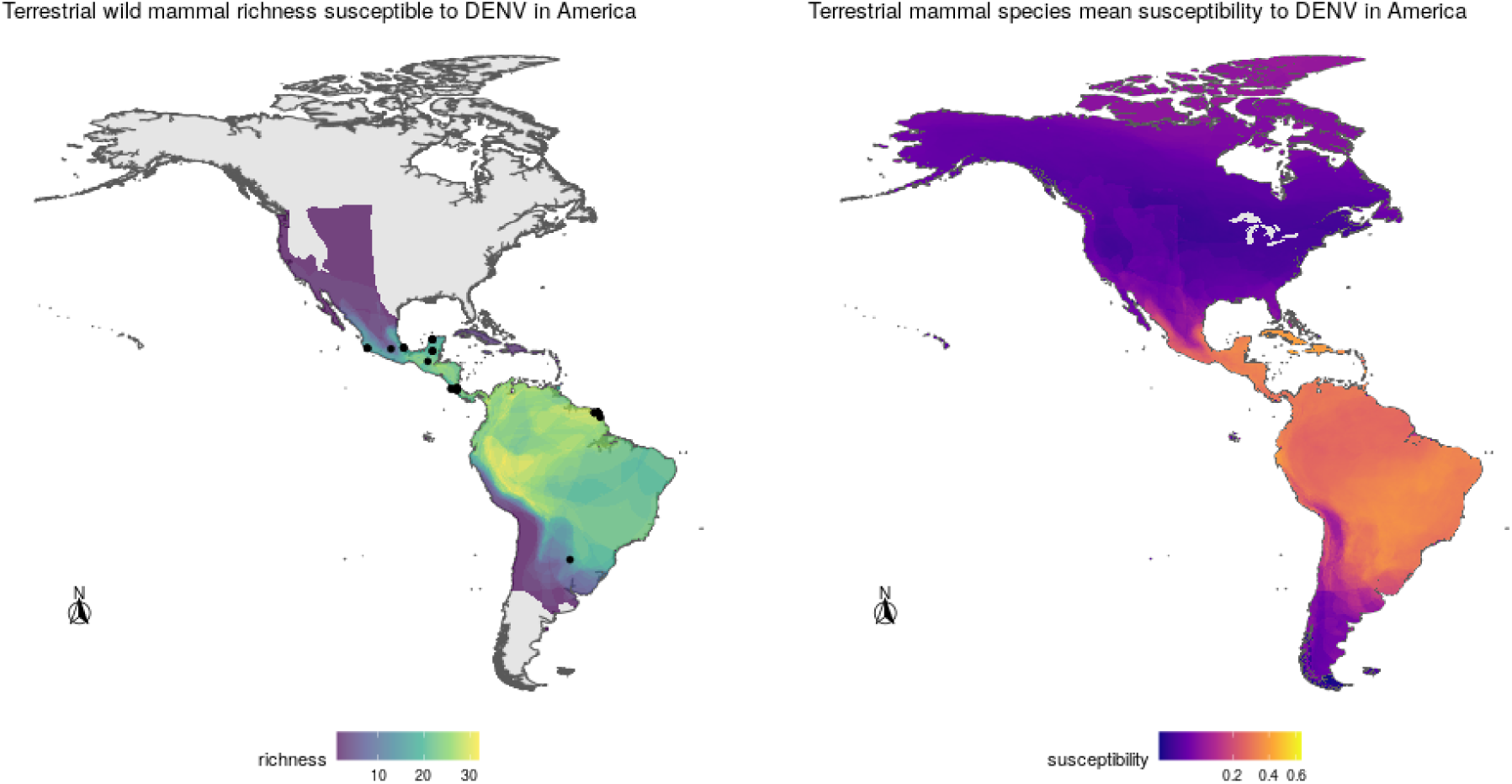
Geographic distribution of richness (left panel) and mean susceptibility to DENV (right panel) of wild mammals in the Americas according to model projections. See Results for details.

**Figure 4.**
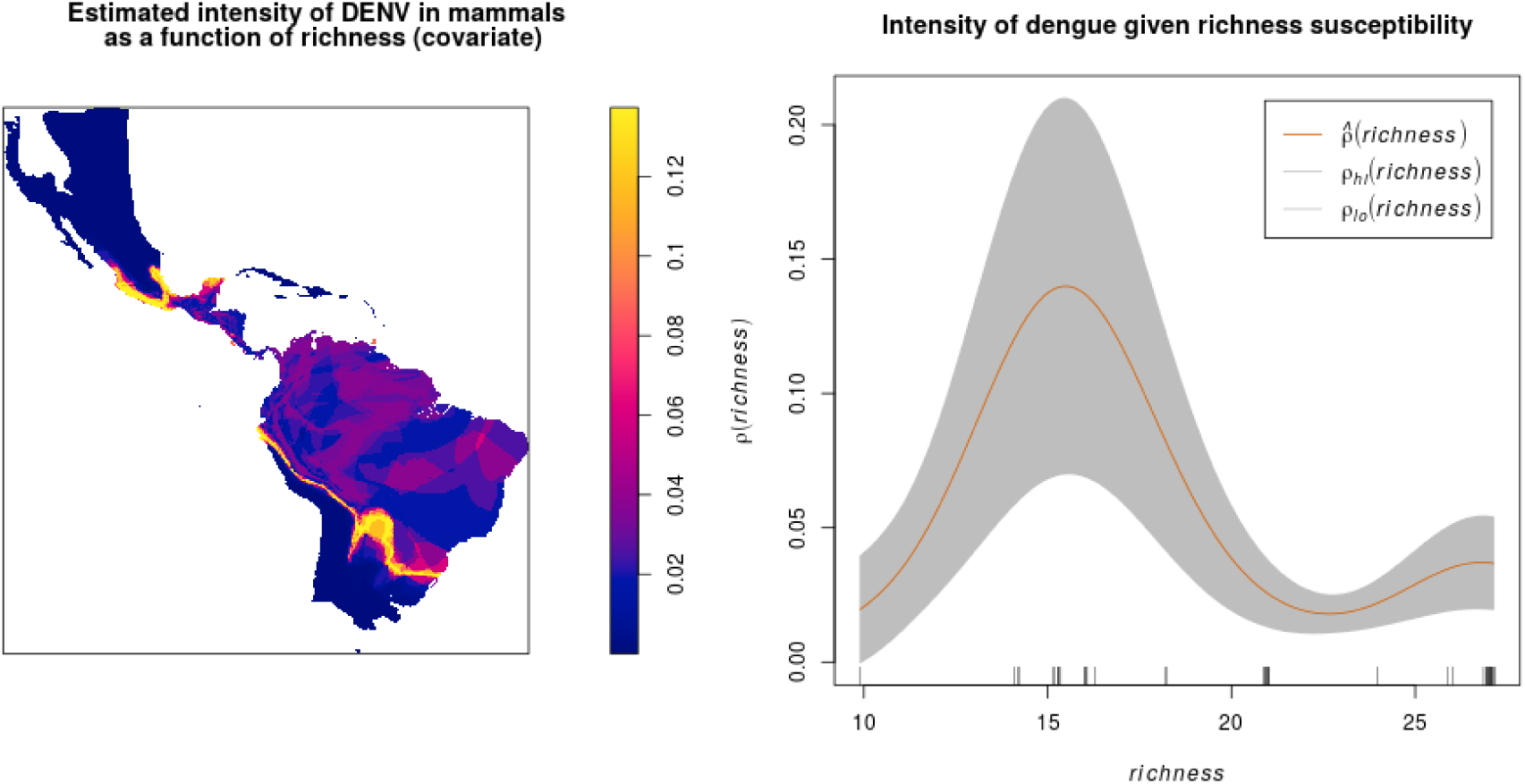
Estimated DENV intensity among American wild mammals given susceptible species richness as a covariate. The plot shows 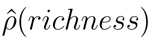 against covariate values of richness, together with 95% confidence bands (between *ρ*_*lo*_ and *ρ*_*hi*_) assuming a non-homogeneous Poisson point process.

**Table 1.** List of top ten highest DENV incidence species according to raw data.

| Order | Family | Species | Incidence |
| --- | --- | --- | --- |
| Chiroptera | Phyllostomidae | *Artibeus jamaicensis* | 66 |
| Primates | Atelidae | *Alouatta caraya* | 48 |
| Chiroptera | Phyllostomidae | *Glossophaga soricina* | 16 |
| Didelphimorphia | Didelphidae | *Marmosa murina* | 15 |
| Chiroptera | Phyllostomidae | *Artibeus planirostris* | 14 |
| Didelphimorphia | Didelphidae | *Didelphis marsupialis* | 12 |
| Chiroptera | Molossidae | *Molossus rufus* | 10 |
| Chiroptera | Phyllostomidae | *Artibeus lituratus* | 9 |
| Chiroptera | Phyllostomidae | *Desmodus rotundus* | 8 |
| Chiroptera | Vespertilionidae | *Myotis nigricans* | 7 |

**Table 2.** List of top ten most DENV susceptible species according to the random forest models.

| Order | Family | Species | Susceptibility |
| --- | --- | --- | --- |
| Chiroptera | Phyllostomidae | *Desmodus rotundus* | 0.909 |
| Chiroptera | Phyllostomidae | *Artibeus libratus* | 0.898 |
| Chiroptera | Phyllostomidae | *Glossophaga soricina* | 0.877 |
| Chiroptera | Phyllostomidae | *Sturnira lilium* | 0.875 |
| Chiroptera | Molossidae | *Eumops perotis* | 0.850 |
| Chiroptera | Molossidae | *Molossus rufus* | 0.845 |
| Chiroptera | Molossidae | *Molossus molossus* | 0.834 |
| Chiroptera | Vespertilionidae | *Myotis nigricans* | 0.821 |
| Chiroptera | Phyllostomidae | *Mimon koepckeae* | 0.821 |
| Chiroptera | Phyllostomidae | *Carollia perspicillata* | 0.814 |

The variable importance for all the models (Figure 2), indicated three most important predictors: the phylogenetic distance between species, the interaction between environmental and geographical distances, and the interaction between environmental and phylogenetic distances (Figure 5). These predictors were followed in importance by environmental distance, while the rest of the predictors and their interactions appeared with lower importance. Although we observed that models had very large variances regarding variable importance, phylogenetic distance remained in the first place of importance.

Regarding the environmental space, mammal species of high incidence and high predicted susceptibility largely overlap in their niche requirements across the Americas (Figure 5). These correspond mostly to warm regions and semiarid to wet areas. Although it is something we had not formally tested (e.g., following ecological niche background similarity tests; Warren et al. 2008), the ecological niches of these two sets of species showed high amount of overlap according to the ellipsoidal models that summarize their ecological niches (Figure 5). The areas with the highest amount of suitable conditions are mostly in tropical America, with extreme values in South America in southern Amazonia, northern Chaco and south of the Eastern Highlands ecoregions, as well as the West Indies and Florida regions, including some parts of Mexico across the Yucatan peninsula and parts of the eastern slope toward the Gulf of Mexico.

**Figure 5.**
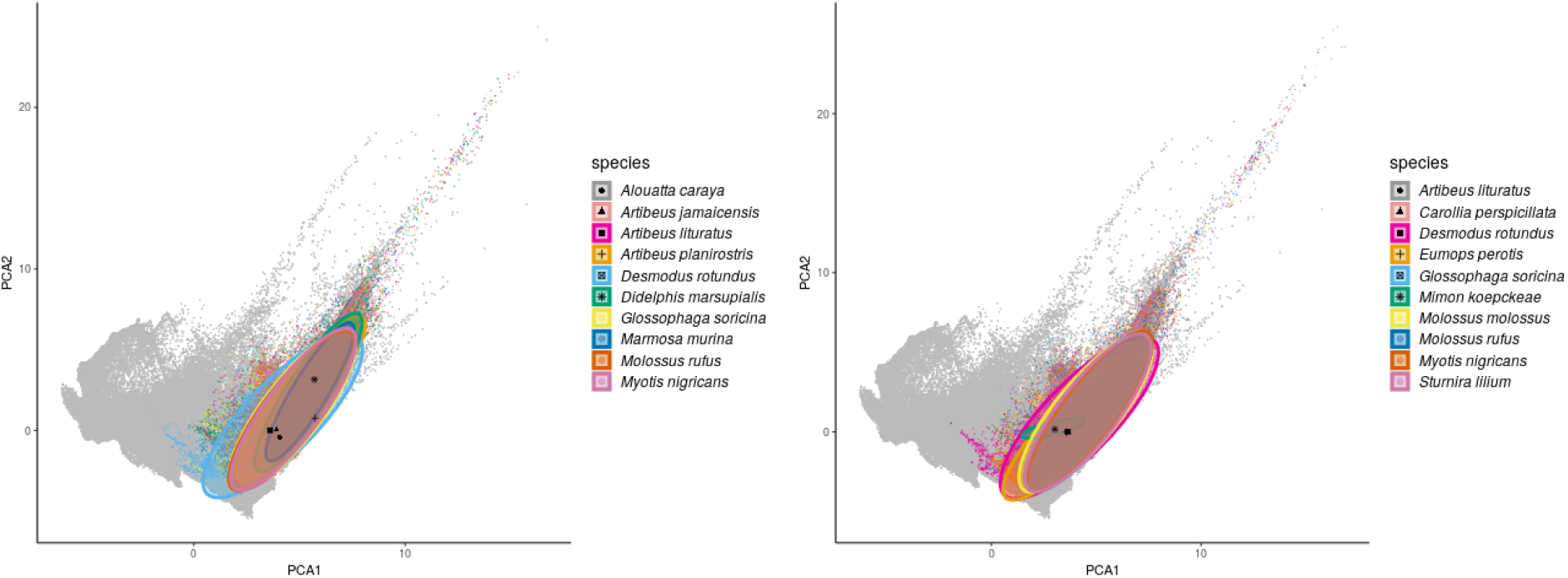
Environmental ellipsoids with centroids. Left panel shows the top ten wild mammal species with highest DENV incidence according to published information. Right panel shows the top ten wild mammal species most susceptible according to random forest models.

**Figure 6.**
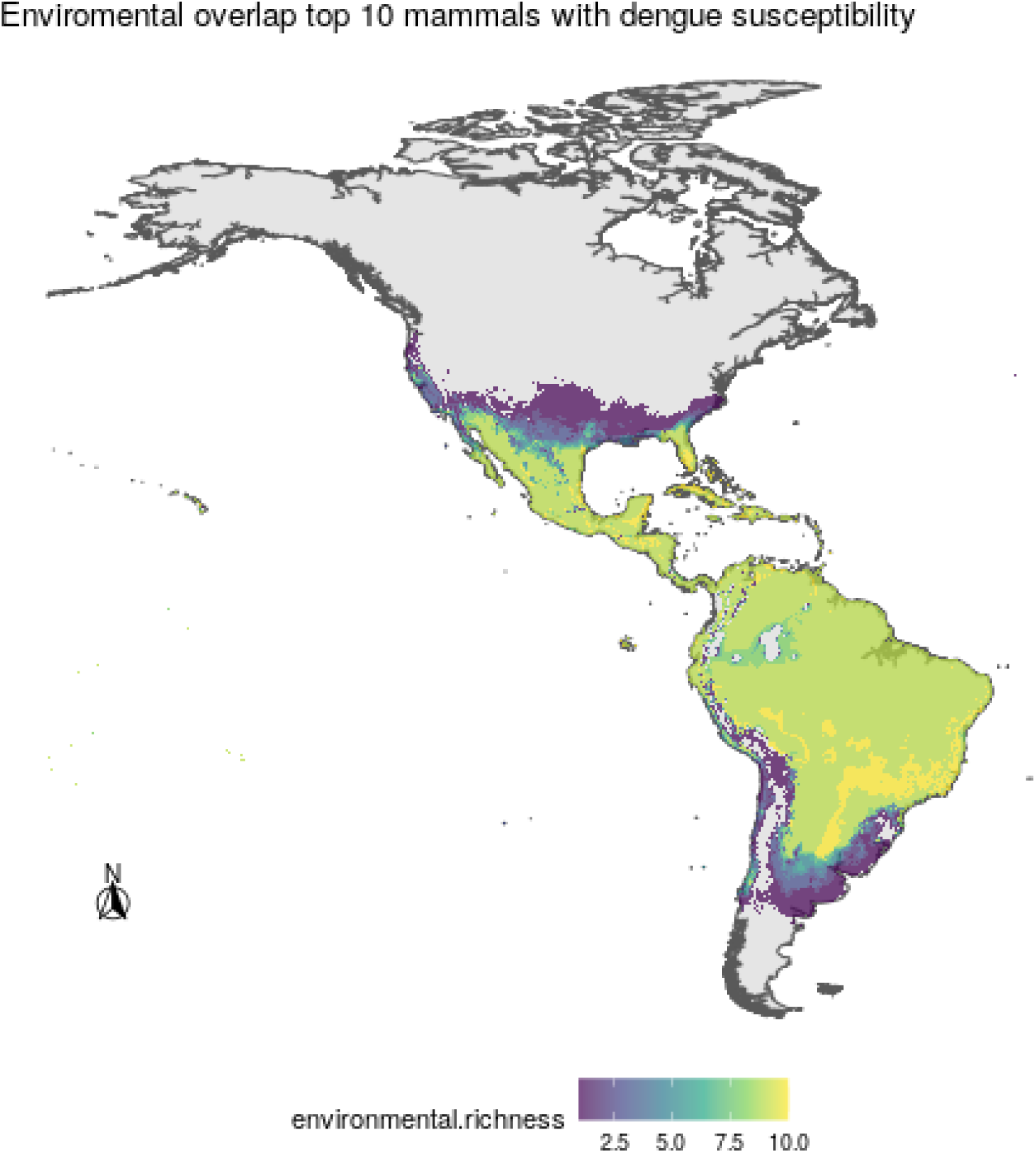
Areas of environmental overlap for the top ten most susceptible wild mammal species to DENV, where higher values indicate more coincidence among species ecological niches.

## 3 DISCUSSION

Researchers have largely overlooked the empirical and theoretical (i.e., model predictions) study of wild reservoir assemblages of DENV. In part this may be due to the great challenge imposed by the difficulty of discovering competent hosts, while we recognize that finding new non-human hosts is also desirable. Here, we show a species-level susceptibility prediction to dengue in wild mammals of the Americas as a function of three biodiversity dimensions (ecological, geographical and phylogenetic spaces). Model predictions coincide with already known geographical distribution patterns for this disease in tropical and subtropical America, where most vulnerable species are bat species of different trophic guilds. Predictions of susceptibility to dengue in wild mammals appears to be highly influenced by phylogenetic distance among species, and also by the environment (i.e., distance between the species’ optima in bioclimatic dimensions) in combination with geographic distance.

To our surprise, most of the empirical and modeling studies of DENV are focused on the urban cycle of transmission (Wearing et al. 2016). Although this is expected given the economic and human health impacts of DENV in urban areas, we suggest that current model projections may fail in the long term if they disregard the role that wild reservoirs (and their phylogenetic, environmental and geographical limiting factors) may play in the ecological transmission of DENV. Wild reservoirs are mostly unknown in the Americas and yet every DENV affecting humans has a sylvatic ancestor (Costa et al. 2012), suggesting that the establishment of a sylvatic cycle in the Americas is only a matter of time. In this sense, our modeling exercise complements all ecological and biogeographic models of previous DENV by providing the perspective from the sylvatic assemblage of potential wild mammal hosts, which may act as an effective ecological reservoir (Haydon et al. 2002). This effort will become even more relevant in the near future, given that anthropogenic activities are bringing closer wild animal assemblages to human populations (Hassell et al. 2017).

The most comprehensive and recent global geographic projection of DENV took into account climatic variables, human population and economic factors, and the distributions of *Ae. aegypti* and *Ae. albopictus* vectors (Messina et al. 2019). Such models continue to predict previously identified tropical and subtropical areas as suitable for DENV, where temperature (68% of variation), precipitation (13%) and relative humidity (6%) are the main environmental variables affecting transmission in that order. Moreover, an estimated average of 3.8 billion people (> 50% of the global population) live in areas suitable for DENV transmission particularly in Asia (Messina et al. 2019). Models predict that for 2050 a large portion of southeastern USA, northern areas of Argentina and higher elevation areas of central Mexico will be suitable for DENV transmission. Some other areas such as East Africa will observe DENV declines due to lower suitability because of hotter and drier conditions (Messina et al. 2019); it is well-known that temperatures higher than 35 degrees Celsius reduce DENV transmission due to reduced mosquito survival (Brady et al. 2014).

The main DENV vector in urban areas is *Ae. aegypti*, which is well-adapted to human dwellings and possesses population characteristics that makes it ideal to maintain DENV infection cycles (e.g., insecticide resistant, desiccation resistance, multiple probing for blood meals on different individuals, capable of using heterogeneous sources for laying eggs avoiding competition and crowded locations; Brady and Hay 2020). Temperature is considered the main climatic factor affecting positively the expansion of DENV across the world, particularly to areas currently free of this virus (e.g., temperate areas of North America and Europe; Brady et al. 2014; Messina et al. 2019). *Aedes albopictus*, one of the most invasive mosquito species worldwide, is suggested to be a competent vector of DENV, and under some situation is even better than *Ae. aegypti*, and what may be limiting its vectorial capacity seems to be more related to its ecology and to its biological attributes (Brady et al. 2014).

The global expansion of the invasive tiger mosquito (*Ae. albopictus*) may aid in DENV geographic expansion. *Aedes albopcitus* has higher tolerance to lower temperatures compared to *Ae. aegypti*, and also inhabits peri-urban and sub-urban areas feeding in a larger array of vertebrate hosts, making it ideal as a bridge vector across the human-domestic-wildlife interface (Wearing et al. 2016). Furthermore, the establishment of DENV in wild reservoirs may aid in the expansion and invasion success of this pathogen. Different species of bats have been positive to different DENV serotypes across the world; in the Americas the four DENV serotypes were detected in French Guiana in a sample that included bats, rodents, and marsupials (Thoisy et al. 2009). In Ecuador and Costa Rica, bats inhabiting urban areas were reported as exposed (seroconverted) to the 1, 2, and 3 DENV serotypes (Platt et al. 2000). In Mexico, DENV-2 serotype has been reported in at least three species of bats in both intact rainforests and disturbed forests in areas of the Yucatan peninsula (Sotomayor-Bonilla et al. 2014); however, in a survey of more than 1900 individuals of different species of rodents and bats on the Pacific slope of Mexico there were no positive results for DENV, which was probably due to the spatiotemporal heterogeneity of transmission (Sotomayor-Bonilla et al. 2018). Furthermore, empirical evidence of non-human primates able to act as reservoirs of epidemic DENV (in particular DENV-2 in an animal breeding facility in the Philippines; Kato et al. 2013) highlights the need to survey potential wild animal reservoirs both in urban (e.g., zoos) and non-urban (e.g., peri-urban forest fragments, agricultural lands) areas, where spillback of urban DENV from humans toward non-human primates or other wild animal can happen (e.g., Teoh et al. 2010). Finally, in a recent analysis within the Flaviviridae, DENV was the only generalist virus capable of infecting several species across the Chiroptera, Rodentia and Didelphimorphia (Sotomayor-Bonilla et al. 2019). Therefore, we suggest that researchers around the world must consider a priority the inclusion of the wild community of potential reservoirs within their DENV surveillance efforts, particularly in areas where the interactions between humans and wild animals is higher (e.g., agricultural lands, peri-urban parks and forests).

Our results confirm that Chiroptera contains a highly susceptible set of species to DENV. In addition, the geographic location of species recorded with DENV from different mammal orders (including those mentioned above; also see table of incidences from this work) confirmed the spatial prediction of our models. This pattern is apparently due to evolutionary and ecological processes as indicated by the importance of phylogenetic and environmental distances in generating these predictions. As past research highlights, phylogenetic distance is one of the most important factors to predict the likelihood of pathogen transmission in hosts of several taxonomic groups not related to mammals (e.g., Gilbert et al. 2012). It is possible that this same process is modulating DENV interaction with mammals. At the same time, however, this should be modulated by species environmental tolerances (i.e., ecological niches) and their spatial context. In fact, this research suggest that highest susceptibility to DENV in American mammals could be constrained not only phylogenetically (as incidence and susceptibility prediction indicate) but also environmentally and geographically. Most species of DENV susceptible mammals appear to share similar environmental requirements (some with wide tolerances, specially in the precipitation axis) and show different degrees of geographic overlap in their distribution areas mostly along the tropical region.

The findings from this biogeographical approach should be put in perspective when DENV-associated reservoirs and vectors are analyzed. Here we highlight the value of combining their phylogenetic, environmental and distributional contexts in order to provide a first effort toward a better understanding of DENVs ecological dynamics in non-urban settings. We hope that these results and methodological approach aids in a solid proposal both to confirm and discover new DENV hosts in the community of wild mammals, while at the same time suggesting what are the main drivers behind DENV transmission.

## AUTHOR CONTRIBUTIONS

All the authors contributed in the data collection, evaluation of the results and writing of the manuscript in its different sections. A.L.R.-F. conducted the analyses.

## ACKNOWLEDGMENTS

A.L.R.-F. received scholarship support from Consejo Nacional de Ciencia y Tecnología (CONACYT; 895979). We recognize the value and effort of the agencies in charge of keeping biodiversity and environmental data available. D.S.-A. was supported by Consejo Nacional de Ciencia y Tecnología (CONACYT, project number Problemas Nacionales 2015-01-1628).

## SUPPLEMENTAL DATA

Supplementary Material should be uploaded separately on submission, if there are Supplementary Figures, please include the caption in the same file as the figure. LaTeX Supplementary Material templates can be found in the Frontiers LaTeX folder.

## DATA AVAILABILITY STATEMENT

The datasets analyzed for this study can be found in the Zenodo repository [https://doi.org/10.5281/zenodo.4018028].

